# Inflammatory Recruitment of Healthy Hematopoietic Stem and Progenitor Cells in the Acute Myeloid Leukemia Niche

**DOI:** 10.1101/2023.11.22.566265

**Authors:** Ding-Wen Chen, Jian-Meng Fan, Julie M. Schrey, Dana V. Mitchell, Seul K. Jung, Stephanie N. Hurwitz, Empar B. Perez, Mauro Muraro, Martin Carroll, Deanne M. Taylor, Peter Kurre

**Affiliations:** Comprehensive Bone Marrow Failure Center, Division of Hematology, Department of Pediatrics, Children’s Hospital of Philadelphia, Philadelphia, PA, USA; Department of Biomedical and Health Informatics, Children’s Hospital of Philadelphia, Philadelphia, PA, USA; Division of Hematology/Oncology, Department of Medicine, University of Pennsylvania, Philadelphia, PA, USA; Single Cell Discoveries, Utrecht, Netherlands; Perelman School of Medicine, University of Pennsylvania, Philadelphia, PA, USA

## Abstract

Inflammation in the bone marrow (BM) microenvironment is a constitutive component of leukemogenesis in acute myeloid leukemia (AML). Current evidence suggests that both leukemic blasts and stroma secrete proinflammatory factors that actively suppress the function of healthy hematopoietic stem and progenitor cells (HSPCs). HSPCs are also cellular components of the innate immune system, and we reasoned that they may actively propagate the inflammation in the leukemic niche. In two separate congenic models of AML we confirm by evaluation of the BM plasma secretome and HSPC-selective single-cell RNA sequencing (scRNA-Seq) that multipotent progenitors and long-lived stem cells adopt inflammatory gene expression programs, even at low BM leukemic burden. In particular, we observe interferon gamma (IFN-γ) pathway activation, along with secretion of its chemokine target, CXCL10. We show that AML-derived nanometer-sized extracellular vesicles (EV^AML^) are sufficient to trigger this inflammatory HSPC response, both *in vitro* and *in vivo*. Altogether, our studies indicate that HSPCs are an unrecognized component of the inflammatory adaptation of the BM by leukemic cells. The pro-inflammatory conversion and long-lived presence of HSPC in the BM along with their regenerative re-expansion during remission may impact clonal selection and disease evolution.

## INTRODUCTION

Acute myeloid leukemia (AML) is a genetically heterogenous disease characterized by clonal expansion of myeloid blasts evolved from hematopoietic stem and progenitor cells (HSPCs) (1). Rationally-designed targeted therapies have led to meaningful improvements in outcome for select AML subgroups, but many patients still succumb to treatment-refractory relapse (2). Recent studies have shown that the inflammatory secretome from AML blasts and bone marrow (BM) stroma plays a role in AML pathogenesis (3, 4). Inflammatory signaling in the niche itself has long been linked to AML progression and hematopoietic dysfunction, (5, 6, 7, 8, 9, 10), and several mechanisms and mediators have been proposed, including NF-κB expression programs in AML blasts (11), and secretion of IL-6 (9) and IL-1β (12). Therefore, while AML blasts (9) and stromal cells (13) in the leukemic niche are known sources of inflammatory factors in the leukemic BM, other cellular contributors remain incompletely understood. Importantly, AML blasts are rapidly eliminated from the leukemic BM with standard treatment, and stroma cells are proportionally few, whereas HSPCs persist and re-expand in the post remission BM. HSPCs sustain lifelong hematopoietic and immune function, adapting during homeostasis and under stress through integration of cell autonomous programs with extrinsic niche signals. The discovery that HSPCs serve as potent sensors and amplifiers of the inflammatory crosstalk in the BM (14) is more recent and implies a potentially durable and potent role in propagating and sustaining an inflammatory state in the compartment (10).

Extracellular vesicles (EVs) generated through several biogenesis pathways are constitutively released from cells, and play a critical role for cell-cell communication in several tissues, including the BM (15). Paracrine and endocrine trafficking of tumor derived EVs contributes to tissue adaptation and facilitates metastatic spread (16). While EV’s function in the context of conventional ligand-receptor based signaling, recent studies from our laboratory and others using xenograft models have shown that purified AML-derived extracellular vesicles (EV^AML^) are by themselves sufficient to suppress hematopoietic progenitors (3, 17), and elicit an unfolded protein response in stromal cells (18). We also showed that long-term hematopoietic stem cells (LT-HSC) escape these suppressive effects and appear to enter a state of reversible metabolic quiescence, effected by EV^AML^ (4, 19).

To assess the inflammatory activation of endogenous, healthy HSPCs in the AML BM and overcome some of the limitations of AML xenografts, including extramedullary (splenic) hematopoiesis, we utilize two congenic murine models of AML. Results show that long-lived HSCs and MPP progenitor populations become inflammatory components of the BM niche, and that their conversion is in part mediated by EV^AML^. Unlike the resolution of acute BM inflammation that follows the elimination of AML blasts by chemotherapy, the adaptation of HSPCs may have long-lasting sequelae beyond early remission.

## RESULTS

### AML elicits inflammation in the BM microenvironment

Inflammation has been implicated in AML pathogenesis (5, 6, 7, 8, 9, 13), but the inflammatory state in the AML niche more broadly, and especially at low leukemic burden, has yet to be systematically interrogated. While our prior studies and those by most other groups rely on AML xenograft approaches, we decided for the current study to investigate inflammation in the C1498 congenic murine model of AML (20, 21, 22, 23, 24). Conceptually, this model enables us to evaluate the leukemic BM in the absence of potential cross-species responses (9), and avoids confounding inflammation from myeloablative conditioning (25). First, to validate the model we intravenously (i.v.) injected C1498 AML cells (CD45.2; 1×10^6^ cells) into healthy non-conditioned C57BL/6J (CD45.1) mice (**Fig. 1A**), with PBS-injected recipient controls. The C1498-engrafted recipients showed a median survival time around 21 days post-injection (**Fig. 1B**), with C1498 AML cells contributing to 24.3±12.4% and 36.1±12.3% of total leukocytes in the peripheral blood (PB) by day 18 and 21, respectively (**Fig. 1C**). PB cell count analysis at these timepoints showed a stable hematocrit, but a declining platelet count starting from day 18 (**Supplementary Fig. 1B-C**). Increased white blood cell counts coincided with elevated circulating leukemic blasts at both timepoints (**Supplementary Fig. 1D**). Necropsy examination of C1498-engrafted mice revealed splenomegaly (**Fig. 1D**), hepatomegaly (**Fig. 1E**) and a significant leukemic burden in the liver (**Fig. 1F**). Interestingly, despite low medullary leukemic burden (4.2±2.2%; **Fig. 1G**), altered HSPC subpopulation frequencies were seen, including relatively reduced short-term hematopoietic stem cells (ST-HSC) and long-term hematopoietic stem cells (LT-HSC), but increased multi-potent progenitor (MPP)-3/4 cells (**Fig. 1H-K**). To delineate the inflammatory state of the leukemic BM at both day 18 and day 21, we utilized a multiplex Luminex platform assay to evaluate the spectrum of secreted factors in the AML niche. Here, levels of CXCL9, CXCL10, and CCL12 were increased over 2-fold in BM extracellular fluid of C1498-injected mice (**Fig. 1L**). The data validate the C1498 model system with low medullary disease burden and recapitulate inflammation in the BM niche reported in patient studies. To assess the translational relevance of the elevated *Cxcl10* observed in our murine model, we also analyzed AML patient BM plasma samples and found elevated levels of secreted *CXCL10* and IL-6 compared to healthy donor BM plasma (**Fig. 1M-N)**. Elevated IL-6 released from blasts in AML BM plasma has been previously reported (9), while elevated *CXCL10* in AML BM plasma is described here for the first time. Together, the data suggest that HSPCs in the C1498 AML niche undergo skewed differentiation with compartmental inflammation that includes the secretion of *Cxcl10*.

**Figure 1.**
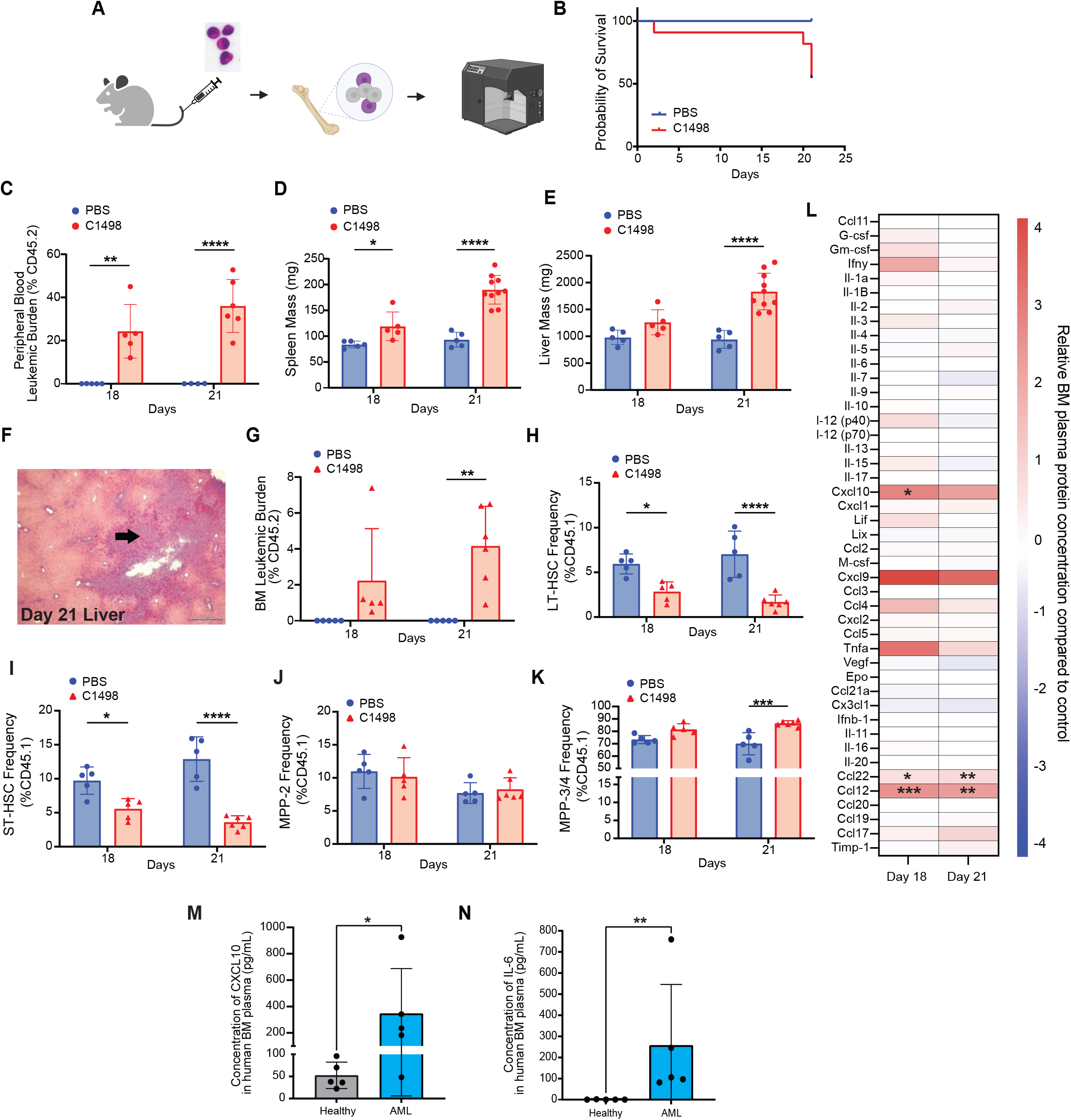
AML elicits compartmental inflammation in BM microenvironment. (A) Schematic of the experimental approach utilizing C1498 AML mouse model (n=5). Endpoint analyses were performed 18- and 21-days post C1498 injection. (B) Survival probability C1498-engrafted mice compared to control. Detectable leukemic burden in PB (C), changes in spleen (D) and liver size (E), and evidence of extramedullary leukemic burden in liver (F) in C1498-engrafted mice. Detectable leukemic burden in BM (G), and analysis of residual HSPC subpopulation: LT-HSC (H), ST-HSC (I), MPP-2 (J), and MPP-3/4 (K) in C1498-engrafted mice. (L) Multiplex analysis of BM plasma in C1498-engrafted mice showed upregulated pro-inflammatory cytokine production compared to control. Validation studies using human BM plasma samples showed elevated pro-inflammatory cytokines, Il-6 (M) and Cxcl10 (N) (n=5)

#### The transcriptome of hematopoietic stem and progenitor cells in the AML niche

To elucidate the potential involvement of HSPCs in inflammatory signaling in the AML niche, we evaluated fluorescent-activated cell sorting (FACS)-sorted HSPCs (CD45.1) from the leukemic niche at animal sacrifice using single-cell RNA sequencing (scRNA-Seq) (**Fig. 2A**). We first analyzed the HSPC (Lin^-^ cKit^+^ Sca1^+^) transcriptome in a pseudo-bulk approach, comparing gene expression between C1498-engrafted (HSPC^C1498^) *versus* PBS-injected (HSPC^PBS^) recipients (**Fig. 2B**). Over-representation analysis (ORA) showed an increase in inflammatory response related pathways (*e.g. Hallmark interferon gamma and interferon alpha pathways*) in HSPC^C1498^ **(Fig. 2C).** Further examination of top differentially expressed genes (**Fig. 2D**) showed that several inflammatory gene targets (*Irf7, Irf8, Stat1, Nfkb1, Nfkb2, Cxcl10)* were significantly upregulated (**Fig. 2E)**. Quantitative real-time PCR (qRT-PCR) confirmed the expression of several inflammation-related gene transcripts in HSPC^C1498^ (**Supplementary Fig. 1E**). Together, scRNA-Seq and qRT-PCR analysis showed that normal HSPCs transcriptionally convert to a pro-inflammatory state in AML BM.

**Figure 2.**
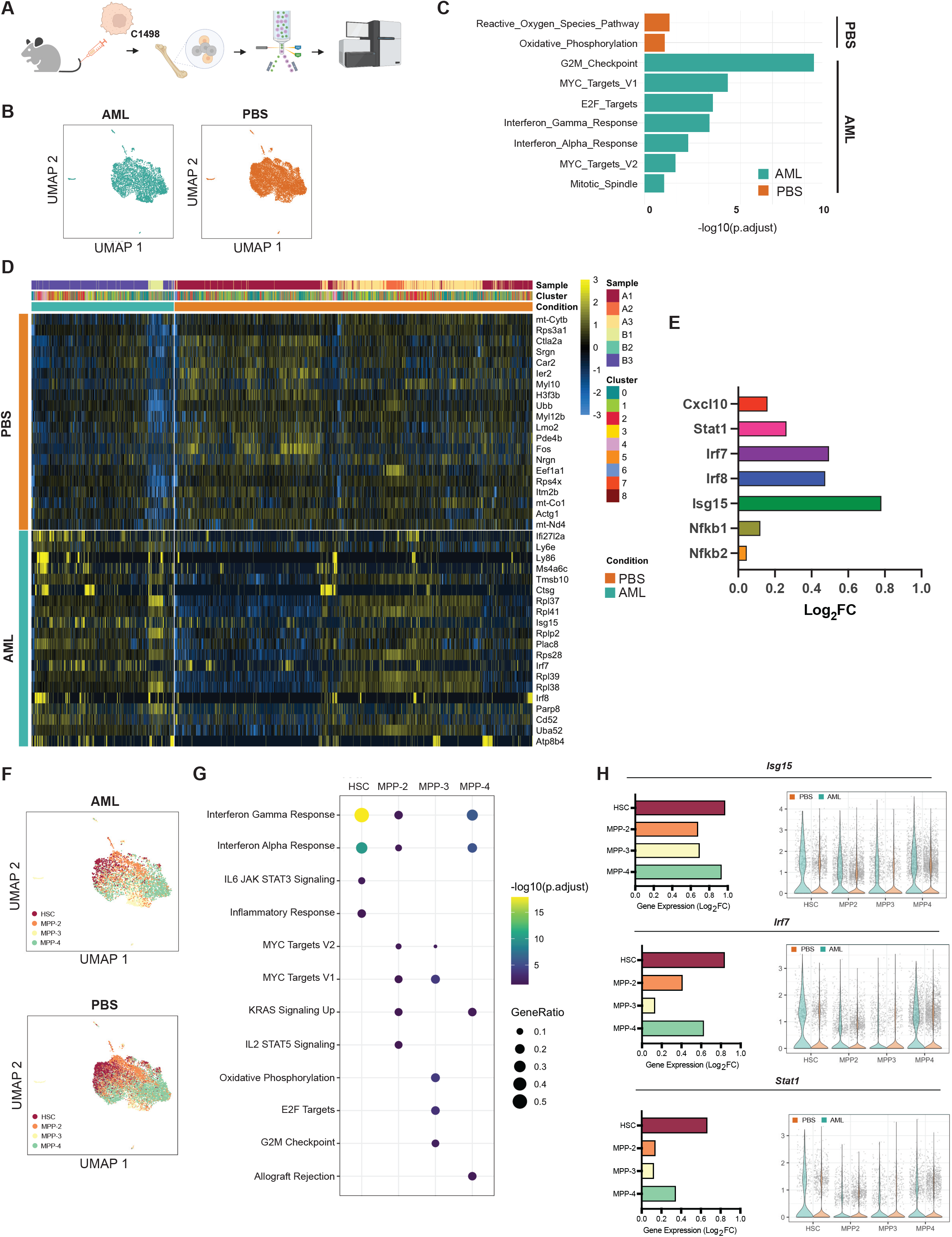
Single-cell transcriptomic analysis reveals hematopoietic stem and progenitor subpopulations exhibit an active inflammatory state in AML. (A) HSPCs from C1498-engrafted AML mice (day 21) were subjected to gene expression analysis (n=3). (B) UMAP of single-cell RNA-sequenced (scRNA-Seq) HSPCs from PBS and AML mice. (C) Gene set enrichment analysis (GSEA) analysis of pathways enriched in HSPCs in AML. (D) Heatmap of top 20 differentially expressed genes by HSPCs in AML and control. (E) Gene expression of selected inflammation-related transcripts in AML. (F) UMAP of HSPC clusters defined into HSC and progenitor subpopulations (MPP-2, MPP-3, MPP-4). (G) GSEA analysis of pathways enriched in HSPC subpopulation in AML. (I) Gene expression of selected inflammation associated genes expressed in HSPC subpopulations in AML.

To understand the potential contribution by specific stem and progenitor populations, we next classified single-cells into HSCs and multipotent progenitors (MPPs; MPP-2, MPP-3, and MPP-4) (**Fig. 2F**) by calculating a module score based on the expression levels of the previously described subpopulation gene signatures (**Supplementary Fig. 2A**) (26). ORA of the list of differentially expressed genes in HSC and MPP clusters showed upregulation of inflammation related pathways (*Hallmark Interferon -Alpha* and *-Gamma* response pathways enriched in HSC, MPP-2, and MPP-4 subpopulations in AML (**Fig. 2G**). Analysis of top differentially expressed genes in the inflammatory response pathways reveal upregulation of several canonical inflammation response genes (*Isg15, Irf7, Stat1)* in HSCs and MPPs (**Fig. 2H; Supplementary Fig. 2D-E)**. Gene ontology (GO) analysis of the genes differentially expressed in healthy HSC from the AML grafted cohort AML shows enrichment of several key processes that are known functional consequences of inflammatory stress such as myeloid differentiation (p-adjusted=1.3×10^-4^) and innate immune processes (p-adjusted=2.4×10^-5^) (**Supplementary Fig. 2C**). Collectively, these results demonstrate for the first time that HSCs and MPPs acquire an active inflammatory state in the AML BM. To conceptually validate the presence of inflammation in HSPCs beyond the C1498 model system, we utilized a more commonly used and translationally relevant MLL-AF9 AML mouse model (27, 28). Typically reliant of retrovirus overexpression, the model system we describe here mimics human disease instead through doxycycline (DOX) mediated regulation of human MLL-AF9 fusion oncogene expression to (29, 30). To evaluate the inflammatory state in HSPCs in the iMLL-AF9 model *in vivo,* we generated chimeric recipients (CD45.2) bearing both iMLL-AF9 BM (CD45.1; iMLL-AF9 leukemic BM) and wildtype BM (CD45.1/2; resembling healthy BM) (**Fig. 3A**). Using this model, we show the rapid emergence of leukemic fraction in the PB over the course of DOX induction (**Figure 3B**). Assessment of BM at day 19 showed high burden of myeloid-restricted MLL-AF9-expressing blasts (**Figure 3C-E; Supplementary Fig 4A-B**), splenomegaly (**Fig 3F**). Gene expression analysis of blasts showed upregulation of key oncogenic transcripts (hMLL-AF9, Meis1, Hoxa9) (**Supplementary Fig. 4C-E).** Importantly, assessment of residual normal HSPCs (CD45.1/2) showed signs of elevated inflammation (**Supplementary Fig. 4F-G**) without significant differences in HSPC subset distribution (**Fig. 3G)**.

**Figure 3.**
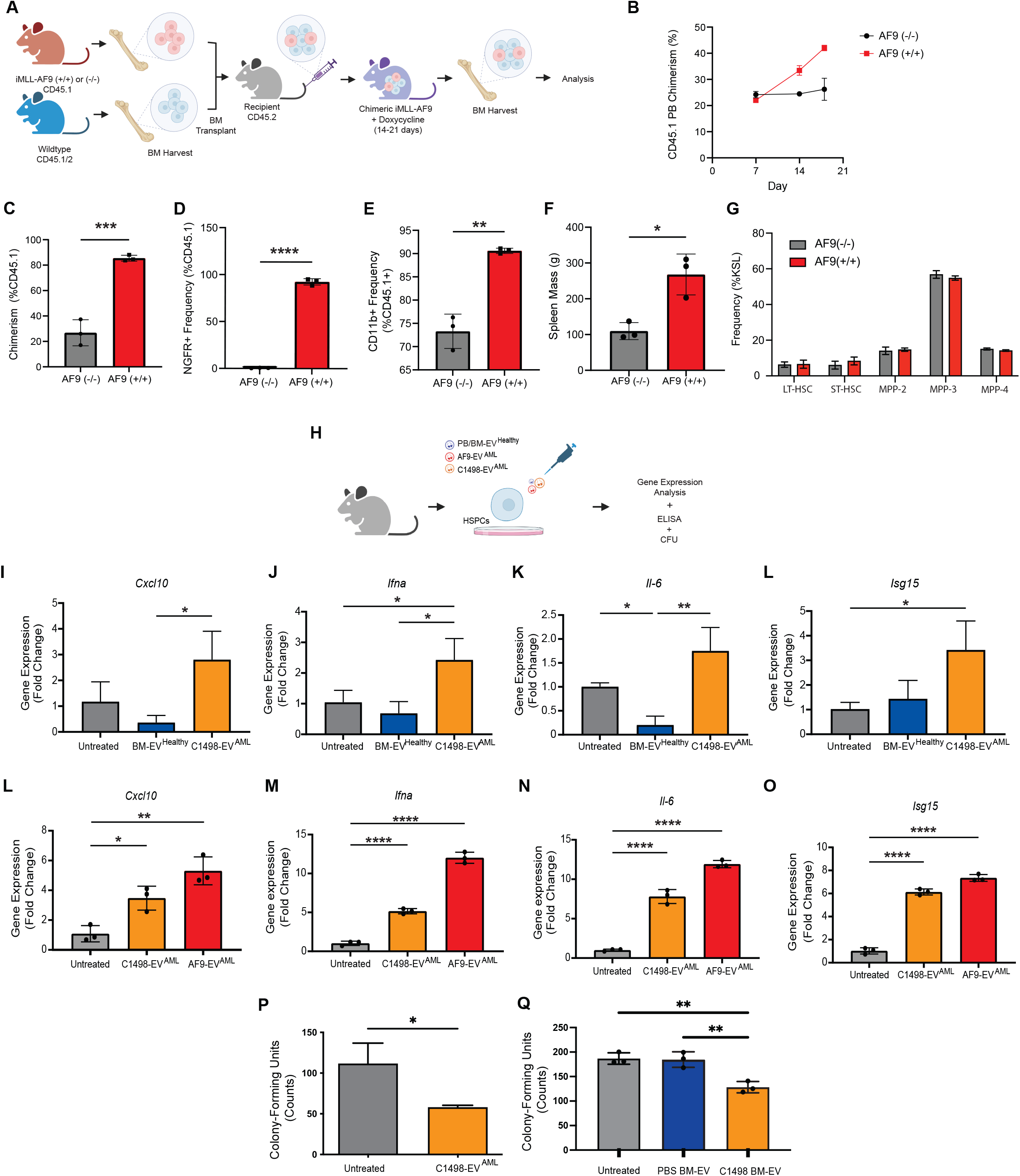
EV^AML^ incites inflammatory responses in HSPCs. (A) A schematic of iMLL-AF9 chimeric mouse model generation. (B) Assessment of leukemic hematopoietic fraction (CD45.1+) in the PB over the course of doxycycline induction. BM was assessed after 18 days of induction (n=3) for leukemic fraction chimerism (C), the frequency of MLL-AF9-expressing cells (based on NGFR expression; cells express MLLAF9-IRES-NGFR cassette) (D) and myeloid cell frequency (E) in the leukemic fraction of the BM. Spleen size (F) and HSPC subpopulation frequency were also assessed (n=3). (H) The role for EV^AML^ in inciting inflammatory responses in HSPCs were carried out by challenging FACS-sorted HSPCs (HSPCs sorted and pool from n=3 mice per experiment) with EV^AML^. Comparison of HSPC gene expression following 2 hours exposure with EV derived from C1498 (C1498-EV^AML^) and healthy BM (BM-EV^Healthy^) (I-L) (n=3). Comparison of HSPC gene expression following exposure with EV^AML^ from both C1498 and iMLL-AF9 blasts (AF9-EV^AML^) (M-P) (n=3). Methylcellulose assay analysis of HSPC colony forming unit counts following 72 hours challenge with either C1498-EV^AML^ (n=3) (Q), or EVs from the BM plasma of C1498-engrafted mice and PBS-injected mice (n=3) (R).

### AML-derived extracellular vesicles incite the inflammatory activation of HSPCs

We were struck by the observation that HSPCs in the C1498 AML niche are converted to an inflammatory phenotype despite low leukemic burden in the BM, and hypothesized that AML-derived extracellular vesicles (EV^AML^) may play a role (3, 4). We first characterized the C1498 AML-derived EVs (C1498-EV^AML^) (**Supplementary Fig. 3A-B**) and confirmed the presence of lipid bilayer morphology vesicles with particle size of 131.0±58.7nm and enriched with known EV protein biomarkers (*Alix, Cd63, and Tsg101*) (**Supplementary Fig. 3C,F**). To determine whether EV^AML^ can directly incite inflammation in HSPCs, we next exposed FACS-sorted healthy HSPCs to C1498-EV^AML^ *in vitro* (**Fig. 3H**), and found transcriptional upregulation of *Cxcl10, Ifn-α, Il-6, and Isg15* (**Fig. 3I-L**). In contrast, HSPC exposed to EVs derived from healthy BM plasma (BM-EV^Healthy^) did not induce upregulation of inflammatory response genes in HSPCs. We further confirmed that EV^AML^ secreted from iMLL-AF9 blasts (AF9-EV^AML^) (**Supplementary Fig. 3D, G**) can also incite upregulation of pro-inflammatory transcripts (*Isg15*, *Il-6*, *Ifn-α*, and *Cxcl10) in naïve HSPC in vitro* (**Fig. 3M-P**). With the observation of elevated *Cxcl10* protein secretion in the BM plasma of C1498-engrafted mice, as well with increased *Cxcl10* gene expression in both C1498- and AF9-EV^AML^-challenged HSPCs *in vitro* (**Fig. 1L; 3I**), our data suggests that HSPCs are actively contributing to *Cxcl10* levels in the AML niche. We excluded *Cxcl10* secretion by C1498 cells, observing no *Cxcl10* protein expression in both C1498-conditioned medium and C1498-EV^AML^ (Supplementary Fig. 5A-B). No detectable *Cxcl10* transcripts were found in C1498-EV^AML^ (**Supplementary Fig. 5C-D)**, nor were there evidence of direct mRNA transfer from EV^AML^ (**Supplementary Fig. 5F-G**), supporting our finding that *Cxcl10* expression is derived from EV^AML^-conditioned HSPCs. IFN gamma (IFN-γ) signaling pathway activation in our scRNA-Seq data, (31), provides a plausible source of stimulation for *Cxcl10*. Functionally, IFN signaling and other inflammatory mediators have been shown to suppress progenitor clonogenicity and HSC stemness (14, 32). To investigate whether inflammatory activation by C1498-EV^AML^ confers hematopoietic suppression in HSPCs, we plated primary HSPCs to methylcellulose colony-forming unit (CFU) assay following C1498-EV^AML^ challenge *ex vivo*. Evaluation of the CFU colony counts confirmed reduced clonogenicity compared after exposure to C1498-EV^AML^ to control (**Fig. 3Q**). Similarly, we observed functional progenitor suppression of HSPCs exposed to EVs derived from BM plasma of C1498-engrafted mice (BM-EV^C1498^). In contrast, HSPCs challenged with EVs derived from BM plasma of healthy controls (**Fig. 3R**) did not exhibit clonogenic suppression. Together, our results indicate that EV^AML^ cause inflammatory activation and suppress HSPC clonogenicity, albeit without linking suppression to one of the inflammatory specific mechanisms.

#### EV^AML^ incite inflammatory responses in HSPCs *in vivo*

To determine whether EV^AML^ can incite inflammatory signaling in HSPCs *in vivo,* we subjected healthy C57BL/6J mice to serial administration (once daily for 3 days) of EV^AML^ (**Figure 4A**). Transcriptomic analysis of HSPCs (KSL) from AF9-EV^AML^-injected mice (HSPC^AF9-EV^) harbored a distinctly different transcriptome profile compared to HSPCs from PBS- and PB-EV-injected recipients (**Fig. 4B-C**). Notably, gene set enrichment analysis (GSEA) revealed enrichment of several inflammatory- and immune response related pathways (**Fig. 4D**). Along with upregulation of several well annotated inflammation related genes (*S100a8*, *S100a9, Sphk1, Mmp14, Clec5a, Sgms2s)* in HSPC^AF9-EV^ (**Fig. 4E**), these results indicate that AF9-EV^AML^ can by themselves induce innate immune responses *in vivo*. Prior studies have reported upregulation of Sca-1 surface protein expression in non-HSPC progenitors (Lin^-^ cKit^+^ Sca1^-^) under inflammatory stress, which can impact conventional HSPC (Lin^-^ cKit^+^ Sca1^+^) immunophenotyping and FACS sorting (33). To exclude this confounder, we compared HSPC subsets assignment using conventional LSK with an alternate immunophenotyping strategy: LSK (Lin^-^ cKit^+^ Sca1^+^; **Supplementary Fig. 6A**) and L86K (Lin^-^ CD86^+^ cKit^+^; **Supplementary Fig. 6B**). These results confirmed that, in contrast with the well described increase in LSK frequency following LPS challenge, there were no significant differences between LSK and L86K frequencies following AF9-EV^AML^ challenge compared to controls (**Supplementary Fig. 6C-I)**. Altogether, our results show AF9-EV^AML^ can elicit the inflammatory conversion in HSPCs.

**Figure 4.**
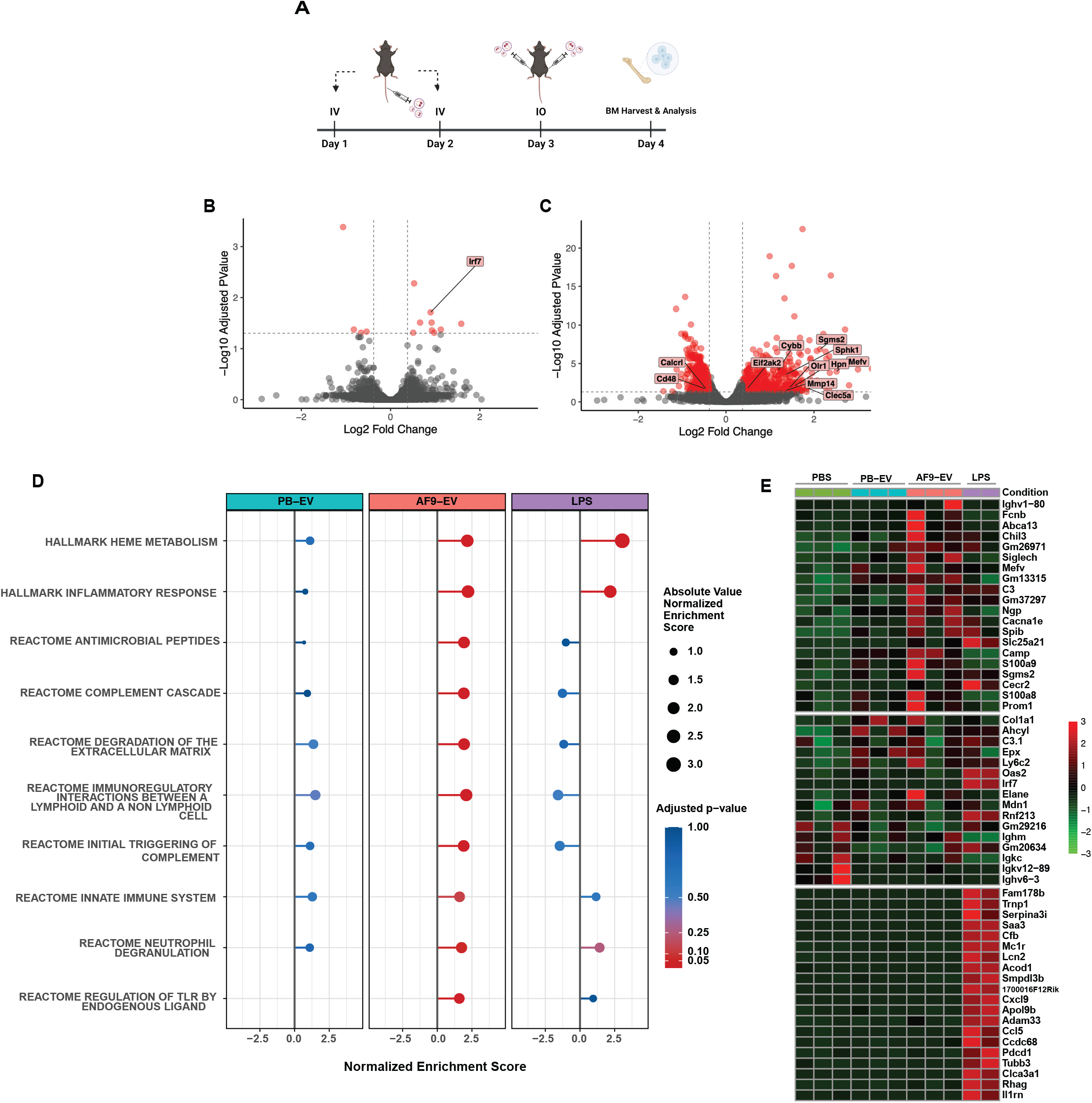
EV^AML^ incites inflammatory responses in HSPCs in vivo. (A) B6 Cd45.1 mice were serially injected with AF9-EV^AML^ (2E9 particles/day) for 3 days (n=3). Mice received either PBS or PB-EV^AML^ (2E9 particles/day) served as negative controls. HSPCs from injected mice were harvested 24 hours the last injection and subjected to RNA-Seq analysis. Volcano plots illustrating differential HSPCs gene expression between: (B) PB-EV and (C) AF9-EV^AML^. (D) GSEA analysis of differential expressed genes in HSPCs from AF9-EV^AML^ showed enriched inflammatory related pathways compared to vehicle control (PBS-injected). (E) Heatmap of top 20 inflammatory targets significantly upregulated in HSPCs from AF9-EV^AML^ injected (top), PB-EV-injected (mid), and LPS-injected (lower panel) mice compared to vehicle control (PBS-injected).

## DISCUSSION

Inflammation has long been associated with AML pathogenesis, accounting for the suppression of healthy HSPCs, leukemia promotion and clonal selection (34). Yet, comprehensive insight into the cellular sources, mediators and mechanisms remains elusive, in part constrained by the reliance on experimental modeling in xenografts. We hypothesized that healthy HSPC are more than simple bystanders and play an active role in the inflammatory leukemic niche. The resulting studies in two congenic murine models of AML now reveal for the first time the extent to which healthy HSPC subpopulations undergo differentiation skewing and a pro-inflammatory conversion in the leukemic niche. We show that this process involves EV^AML^ crosstalk that activates interferon pathways and upregulates expression of the pro-inflammatory chemokine *Cxcl10*, among others, in healthy HSPCs.

Prior studies of AML link inflammatory cytokine secretion to AML blasts and BM stromal components (6, 12). Unlike reports in myeloproliferative neoplasms, the role of residual healthy hematopoietic cells in AML and their contribution to the inflammatory secretome has received relatively little attention (35, 36). Here, we dissected the role of immunophenotypic HSPC and demonstrate that the single cell transcriptome aligns with the BM plasma secretome even at low leukemic burden. Work in both C1498 and iMLL-AF9 AML models consistently demonstrates that HSPCs undergo an inflammatory conversion in the AML niche that predominantly involves interferon signaling pathways (37). GSEA and GO analysis of HSCs in AML reveal pathway enrichment for myeloid differentiation and innate immune response processes, consistent with reports of models of experimental LPS-induced inflammation in HSPC (38). Intriguingly, immunophenotypically defined HSCs show robust *Cxcl9* and *Cxcl10* gene expression, when neither is known to be secreted by AML blasts and both are known to be upregulated by IFN-γ pathway activation.

Our studies further revise the view of HSPCs as passive bystanders that are physically displaced from niche occupancy by encroaching AML blasts. Several groups showed that active crosstalk by leukemic cells at low tumor burden can elicit progenitor suppression while preserving long-lived HSCs in the BM (39, 40, 41, 42, 43). However, while these changes are reversible, adaptation to sustained inflammatory stress may impose durable changes in HSC with potential generation of preleukemic clones, (41, 44), or as a source of selective pressure that promotes clonal amplification (45, 46). For example, clonal hematopoiesis with indeterminate potential (CHIP), more commonly in AML patients, is one of the conditions closely associated with inflammation, wherein clonal expansion compromises long-term HSC function (47). Inflammation significantly alters fate and function of HSC through regulation of survival factors, myelopoiesis and immunosuppressive response (38), but clonal restriction through selective expansion itself can lead to systemic inflammatory sequelae (48).

Adding HSPC to the list of cellular sources of the inflammatory secretome, the current observations expand our understanding of the AML BM as a self-reinforcing inflammatory niche and potential sanctuary for disease persistence (12). HSPC secreted factors, such as *CXCL10* have not been evaluated in the setting of drug resistance in AML, but have been suggested to promote chemoresistance (49) and blast migration to the BM in an acute lymphoblastic leukemia model (50). Other studies showed that *CXCL10* suppresses colony formation by human hematopoietic progenitors (51), and acts on T-cell generation and trafficking (52).

The translational significance of an inflammation-primed HSPC compartment also informs our understanding of the delayed hematopoietic recovery of cytopenic patients (14, 53). Indeed, a recent study in pediatric and adult AML patients showed that inflammation is present at any stage of AML, independent of tumor burden (9), and denoting worse overall survival (54). This has led to the development of a new scoring system that incorporates into an outcome risk score (54). A corroborating study reports the benefits of suppressing inflammation in AML patients with hyperleukocytosis that received chemotherapy with or without dexamethasone (55). Moreover, HSPCs may serve as long-lived cytokine producers, potentially contributing to drug resistance and relapse in AML (56, 57, 58).

While our study provides unambiguous evidence for inflammatory adaptation of HSPC in the leukemic BM, we cannot comment on the long-term consequences and durability. We also cannot comment on the exclusivity by which EV elicit inflammation, as the data do not fully exclude all components of the non-EV secretome from the observed effects. It should also be acknowledged that It is possible that inflammation may vary depending on AML subtype and age of the patient.

Altogether, our report for the first time provides evidence that healthy HSPC contribute to the compartmental inflammation in the AML niche. Such a conversion can be elicited by EV trafficking, leaving HSPCs as long-lived hubs of inflammatory activity with a potentially profound impact subsequent response to infection or re-emergence of disease.

## METHODS

### Mice

Wildtype C57BL/6J-CD45.1 (B6.SJL-Ptprc^a^ Pepc^b^/BoyJ) mice were purchased from The Jackson Laboratory. iMLL-AF9 ((NOD.Cg-Kit ^W-41J^ Tyr ^+^ Prkdc^scid^ Il2rg^tm1Wjl^/ThomJ) were generously gifted by Dr. Shangqin Guo (Yale School of Medicine, Yale University). B6 CD45.1/2 were generated by crossing WT C57BL/6J with B6 CD45.1 mice for experiments. All animal experiments were conducted in accordance with Institutional Animal Care and User Committee (IACUC) protocol at the Children’s Hospital of Philadelphia

### Cell Culture

C1498 cells (also known as TiB-49) were obtained from Dan Marks (Oregon Health & Science University). C1498 cells were grown in Dulbecco’s Modified Eagle Medium (DMEM) (Corning, Corning, NY, USA) supplemented with 10% fetal bovine serum and penicillin/streptomycin (100U/mL) and cultured cells in humidified incubator chamber at 37°C/5% CO2. HSPCs (Lin^-^ cKit^+^ Sca1^+^) were FACS-sorted and cultured in liquid culture using StemSpan (Stem Cell Technologies, Vancouver, Canada) supplemented with Il-3 (10ng/mL; Peprotech,Cranbury, NJ, USA), Il-6 (10ng/mL; Peprotech), Scf (50ng/mL; Peprotech), and PenStrep (1%). iMLL-AF9 blasts were cultured ex v*ivo* in StemSpan supplemented with Tpo (50ng/mL; Peprotech), Flt3-l (50ng/mL; Peprotech), Scf (100ng/mL; Peprotech), and DOX (2μg/L; Sigma; St. Louis, MO, USA).

### *In vivo* procedures

*<u>iMLL-AF9 chimeric mice</u>* were generated by co-transplanting BM harvested from homozygous iMLL-AF9 mice (CD45.1; 5E5 cells) and WT B6 mice (CD45.1/2; 5E5 cells) into lethally irradiated C57BL/6J (CD45.2) recipients. AML leukemogenesis was initiated via doxycycline (DOX) water feed (supplemented with 1g/L DOX and 10g/L sucrose) 4-8 weeks post BM transplantation. *<u>Serial in vivo EV injection</u> <u>experiments</u>* were performed by injecting either AF9-EV^AML^ (1-2E9 particles per injection) or PB-EV (1-2E9 particles per injection) into B6 CD45.1 mice every 24 hours for 3 days via intravenous (for the first two injections) and intraosseous (third injection). Either PBS or lipopolysaccharides (LPS; 3mg/kg; 1 dose intraperitoneal) were injected as controls.

### FACS cell sorting or analysis

Cells were sorted and analyzed using either FACSAria Fusion cell sorter or FACS Canto II (Becton Dickinson, Franklin Lakes, NJ, USA). Lineage-negative cells were enriched using EasySep Mouse Hematopoietic Progenitor Cell Isolation Kit (Stem Cell Technologies) according to manufacturer’s protocol prior to sorting. Cells are stained with appropriate mouse antibodies (Supplementary Table 2).

### Extracellular vesicles isolation and characterization

Extracellular vesicles were harvested via differential centrifugation methods involving: 400 x g for 10 mins, 2,000 x g for 20 mins, 10,000 x g for 20 mins, and 100,000 x g for 2 hours. Ultracentrifugation were performed using either Optima L-60 or L-80 instrument with either SW-28, SW-40, or SW-55-Ti rotors (Beckman Coulter, Brea, CA, USA). Pelleted EVs were resuspended in PBS. EVs were quantified utilizing tunable resistance pulse sensing method using qNano Gold (Izon, Christchurch, New Zealand), with either NP150 or NP200 nanopores.

### Transcriptional gene expression assays

RNA from FACS-sorted HSPCs were extracted using RNEasy Micro Plus Kit (Qiagen, Hilden, Germany) according to manufacturer’s protocol. *<u>qRT-PCR:</u>* Reverse transcription were performed using SuperScript IV RT (Thermo Fisher Scientific, Waltham, MA, USA) and qRT-PCR using appropriate primers (Supplementary Table 1). *<u>RNA-Seq</u>:* Libraries were generated using NEX Rapid Dir RNA-Seq 2.0 Kit (Perkin Elmer) and sequenced using NovaSeq (Illumina, San Diego, CA, USA). RNA-seq data were preprocessed using the nf-core/RNAseq pipeline version 3.9. In brief, FastQC version 0.11.9 was used to assess the quality of 100 bp paired-end reads. Trim Galore! version 0.6.7 was then used to trim adapters and low quality bases from raw reads. Reads were aligned to GRCm38 (mm10) using STAR version 2.7.10a and quantified using RSEM version 1.3.1. Duplicates were removed using picard MarkDuplicates version 2.27.4-snapshot. Following preprocessing, data were filtered using the R package WGCNA version 1.69 to remove genes with many missing entries or zero variance. Differential expression between pairs of groups was estimated using DESeq2 version 1.38.3. Genes were considered to have significant differential expression if there was an adjusted p-value of less than 0.05. Pathway enrichment analysis was performed using genes with significant differential expression between each group pairing ranked by their log2foldchange using the R package fGSEA version 1.24.0. *<u>scRNA-</u> <u>Seq</u>:* Gel-in-beads and libraries were generated using Chromium Next GEM Single Cell 3’ Reagent Kits v3.1 (10x Genomics) and sequenced using NovaSeq (Illumina). Cell Ranger 7.0.1. was used to map the sequencing data. The raw sequencing data (FASTQ files) was mapped to the mouse reference genome 2020-A (GRCm38, GENCODE vM23/Ensembl 98). Data from all samples were loaded in R (version 4.2.2) and processed using the Seurat package (version 4.3.0 – Hao et al. Cell 2021). Cells with low UMI counts were filtered out. The filtering thresholds were determined independently for each library to exclude outliers (A1: 1800 UMIs/cell; A2: 3000 UMIs/cell; A3: 1800 UMIs/cell; B1: 2000 UMIs/cell; B3: 2000 UMIs/cell). Cells with more than 10% mitochondrial genes were also removed. Cells from library B2 were excluded due to the low quality of the library. Cells were annotated using gene signatures for HSPCs identified reported by Pietras et al (26). The gene signature for each population was defined by the top 100 genes ranked by the Significance Analysis of Microarrays (SAM) scores. We calculated a module score for each cell based on the average expression levels of every gene signature, subtracted by the aggregated expression of randomly selected control genes. Each cell was assigned to the population with the highest positive score. Cells with only negative scores were annotated as “Unclassified”. Differentially expressed genes between conditions (AML vs PBS) per cell type were calculated using the Wilcoxon rank sum test. Over-representation analysis of the differentially expressed genes was performed using the enricher function in the clusterProfiler package (version 4.6.2) with the collections “MH: hallmark gene sets”, “M2: curated gene sets”, “M5: ontology gene sets” and “M8: cell type signature gene sets” from the Mouse Molecular Signatures Database.

## Human bone marrow plasma samples

Human BM plasma samples from AML patients and healthy donors were obtained from Dr. Martin Carroll and Dr. Nicolas Skuli from the Stem Cell and Xenograft Core (Perelman School of Medicine, University of Pennsylvania).

## Graphical illustrations

Illustrations were generated utilizing a licensed application BioRender with rights to use in publications.

## Protein Assays

BM plasma proteins were assayed using either enzyme-linked immunosorbent assay (Thermo Fisher Scientific) or mouse cytokine/chemokine 44-Plex Discovery Assay performed by Eve Technologies (Calgary, Alberta, Canada).

## Statistical Analysis

*Statistical significances are calculated using T-Test or ANOVA where appropriate*.

## Acknowledgements

We thank Flow Cytometry Core (CHOP), Center of Applied Genomic Core (CHOP), Bioinformatics Core (CHOP), and Electron Microscopy Resources Core (Penn) for their instrumentation and assay support. We also thank Single Cell Discoveries for their help with data analysis. We also thank Alex’s Lemonade Stand Foundation and Cure4Cam Childhood Cancer Foundation for their generous funding support.

## Statement of Contributions

*Conceptualization: D.W.C., P.K.*

*Methodology Development: D.W.C., P.K*.

*Data Acquisition: D.W.C., J-M. F., J.S., S.K.J., S.N.H*.

*Data Analysis: D.W.C., J-M. F., D.V.M., D.T. E.P., M.M*.

*Manuscript Writing and Editing: D.W.C., J.S., S.N.H., P.K*.

*Funding: D.W.C., P.K*.

## Data Availability Statement

Raw sequencing data are deposited into Gene Expression Omnibus (GEO) database and accessible by the public. GEO Number:

## Figure Legends

***Supplementary Figure 1 – Additional characterization data relating to C1498 model***

***Supplementary Figure 2 – Annotated HSPC subpopulation (HSC, MPP-2, MPP-3, and MPP, HSPC differentially expressed genes, and HSC gene ontology analysis*.**

***Supplementary Figure 3 – Characterization of EVs derived from C1498 cells, iMLL-AF9 blasts, and healthy peripheral blood*.**

***Supplementary Figure 4 – Characterization of iMLL-AF9 mouse model in vivo*.**

***Supplementary Figure 5 – Evaluation of C1498-EV^AML^ cargo and inflammation***

***Supplementary Figure 6* – Comparison of LSK and L86K HSPC frequency following EV injection challenge *in vivo*.**

***Supplementary Table* 1 – Sequences for quantitative PCR analysis primers**

**Supplementary Table 2 – List of antibodies used for immunofluorescence staining**

