## Supplemental Figures and Tables for "Inflammatory Recruitment of Healthy Hematopoietic Stem and Progenitor Cells in the Acute Myeloid Leukemia Niche"

### Supplementary Fig 1

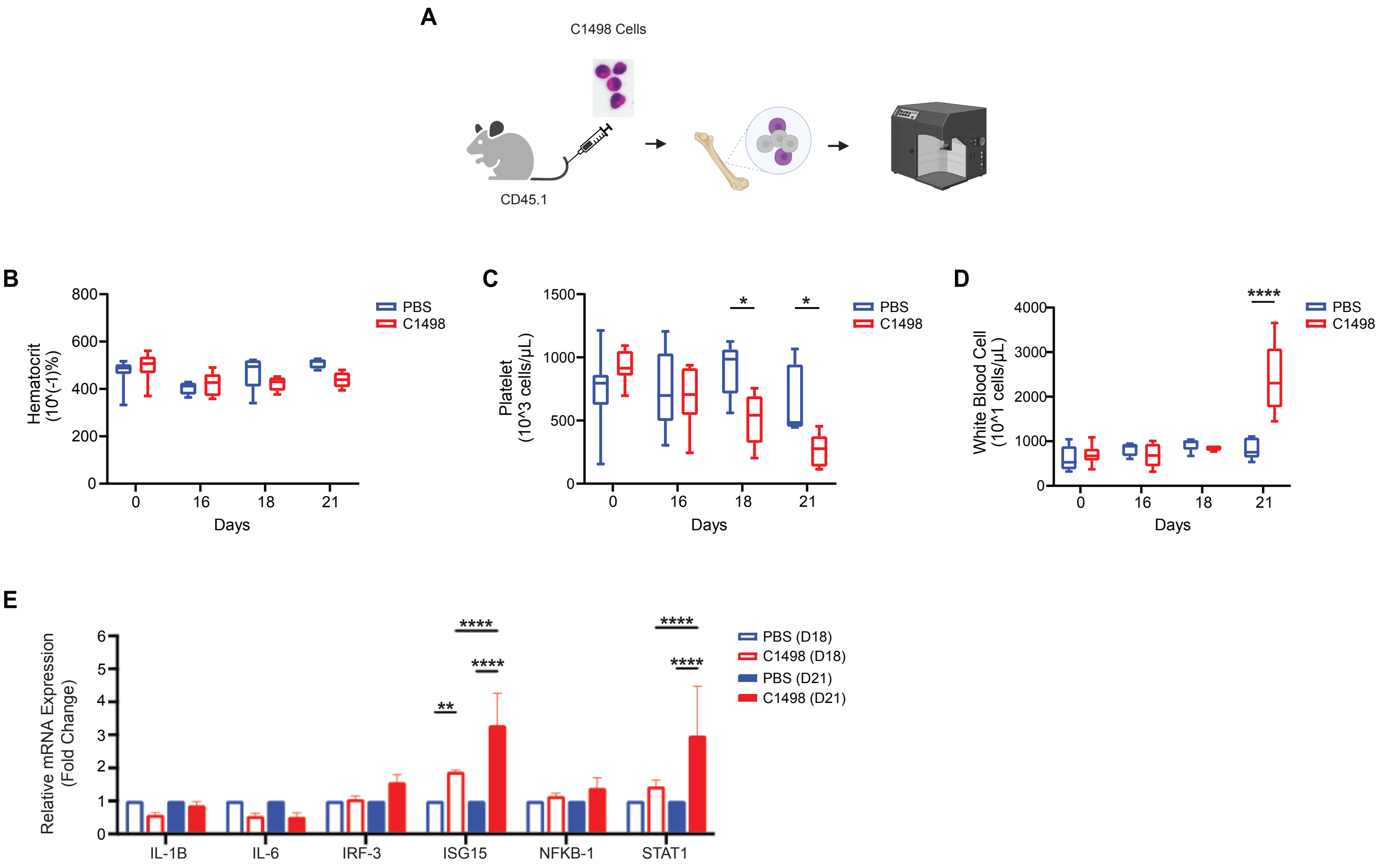


Supplementary Figure 1 – Additional characterization data relating to C1498 model. Complete blood count analysing showing hematocrit (B), platelet (C), white blood cell (D) counts. Quantiative real-time PCR analysis of HSPCs (Lin- cKit+ Sca1+) gene expression 18 and 21 days post C1498 engraftment.

### Supplementary Fig 2

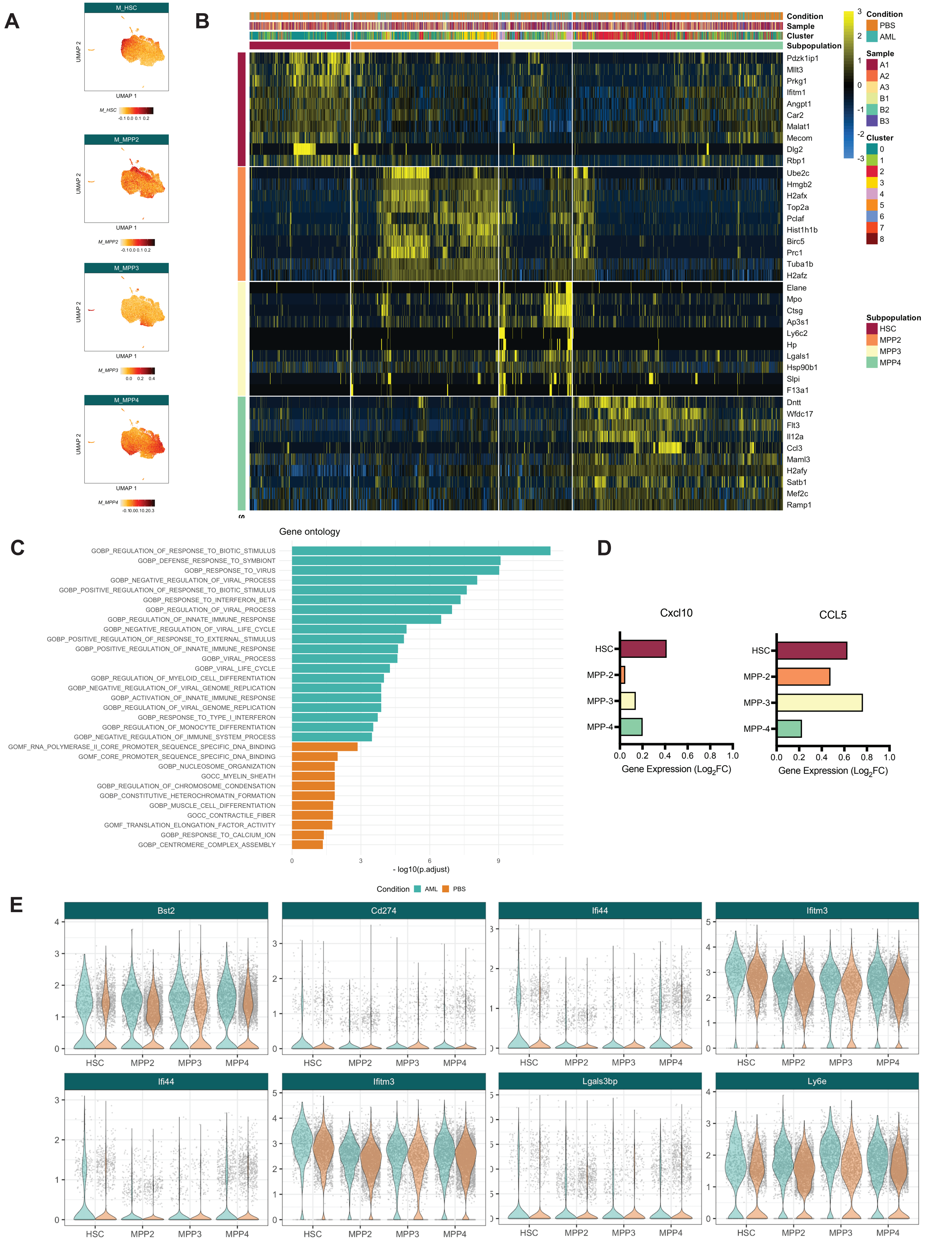


Supplementary Figure 2 – (A) UMAP showing module score of HSC, MPP-2, MPP-3, and MPP-4 gene signatures. (B) Top differentially expressed genes between HSC, MPP-2, MPP-3, and MPP-4 subpopulations. (C) Gene ontology (GO) analysis of enriched processes in HSCs. (D) Selected inflammatory cytokine encoding genes (E) and leading edge genes from Interferon Response Pathways upregulated in HSC and MPPs in AML.

### Supplementary Fig 3

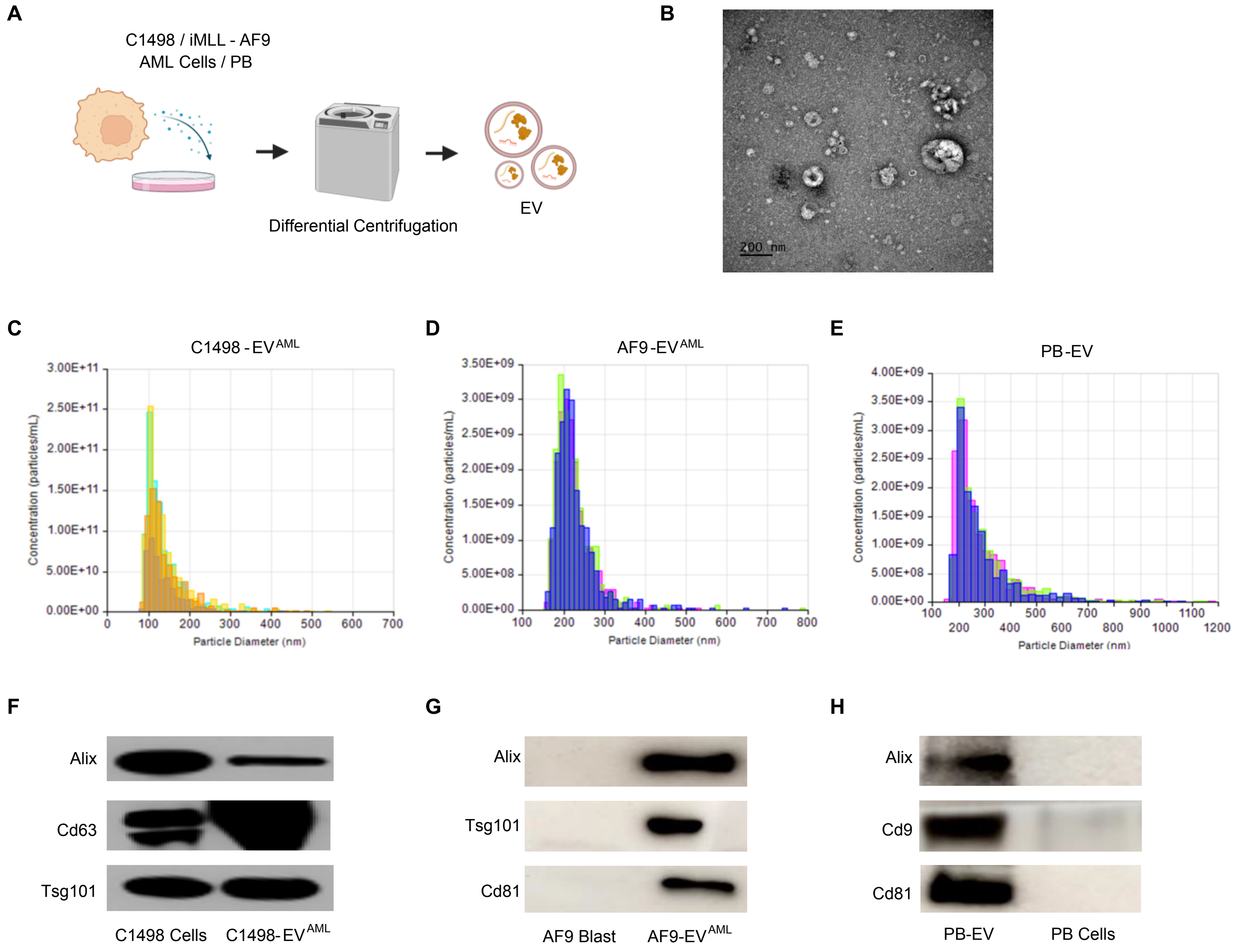


Supplementary Figure 3 – Extracellular vesicle harvest schematic (A). Transmission electron microscopy image of C1498-EV$^{AML}$ (B). EV particle quantitation and biomarker characterization for C1498-EV$^{AML}$(C,F), AF9-EV$^{AML}$(D,G), PB-EV$^{Healthy}$ (E,H),

### Supplementary Figure 4

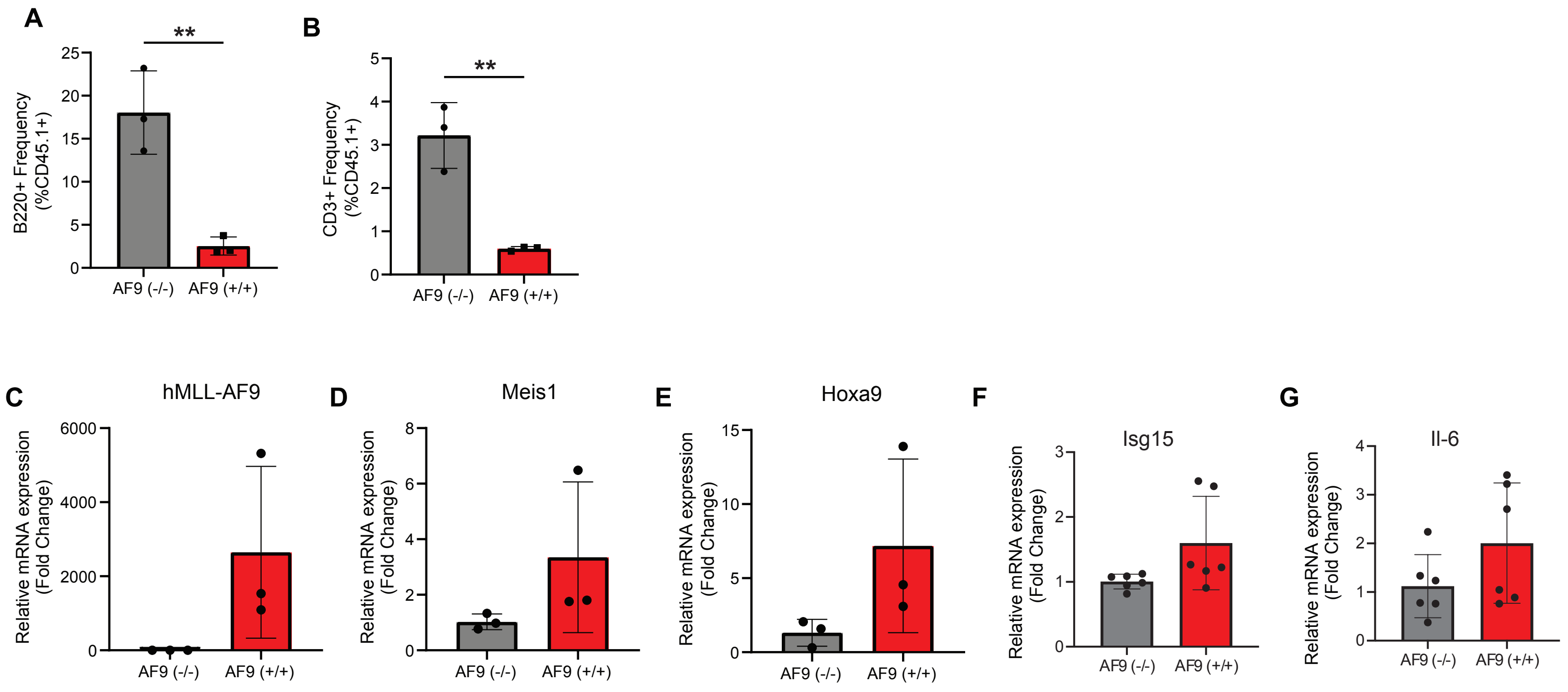


Supplementary Figure 4 – BM B220+ CD3+ subsets in iMLL-AF9-expressing BM cells (A-B). Gene expression analysis of AF9 blasts (CD45.1+; C-D) and normal HSPCs (CD45.1/2+; F-G).

### Supplementary Fig 5

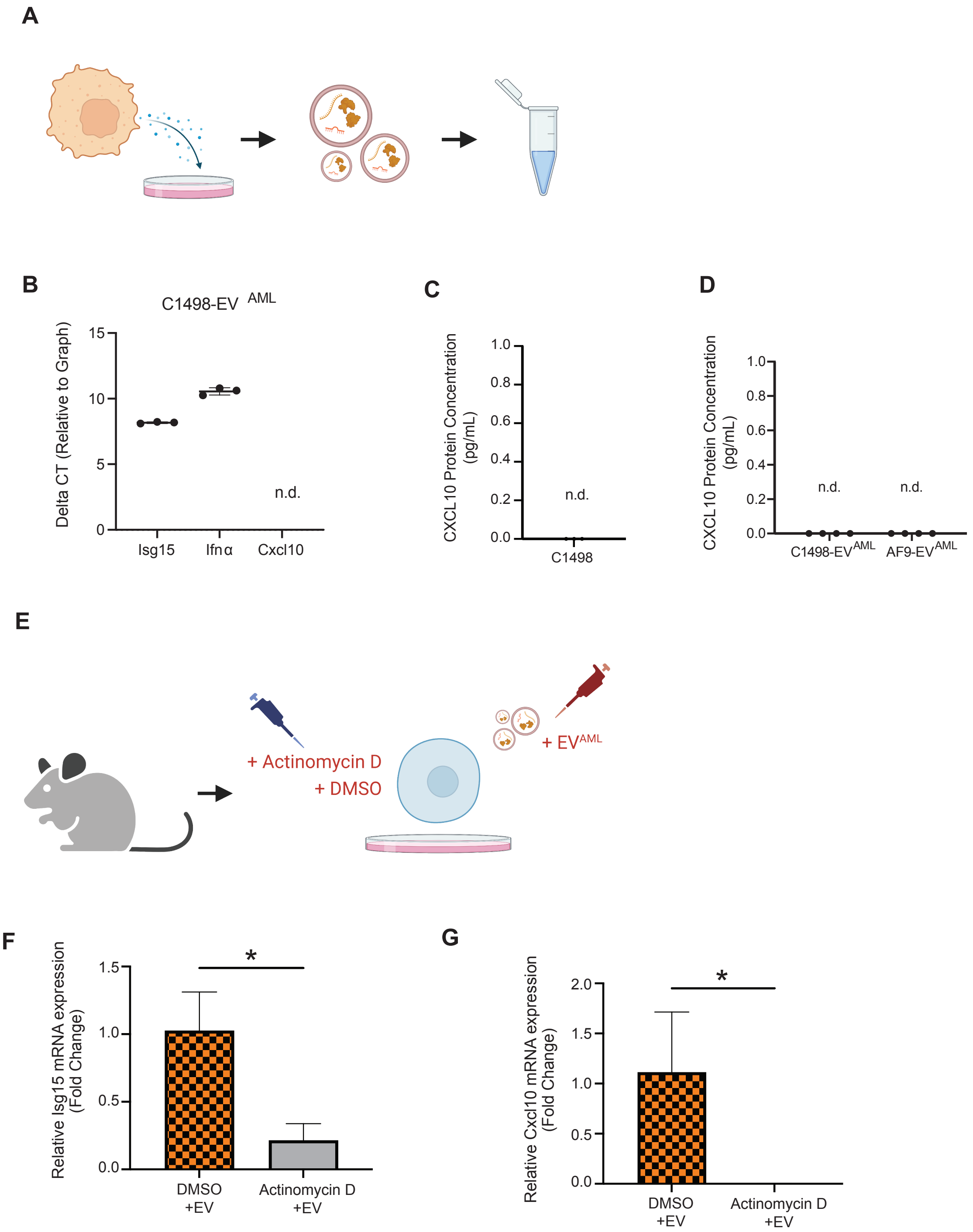


Supplementary Figure 5 - Schematic of RNA analysis in EVs (A) and relative delta CT of target RNA compared to GAPDH (B). Cxcl10 proteins were not detected (n.d.) in C1498-conditioned medium (C) and $EV^{AML}$ (D). Schematic of Actinomycin D treatment experiment (E), and the gene expression analysis of HSPCs following EV challenge, with or without Actinomycin D pretreatment (F-G).

### Supplementary Fig 6

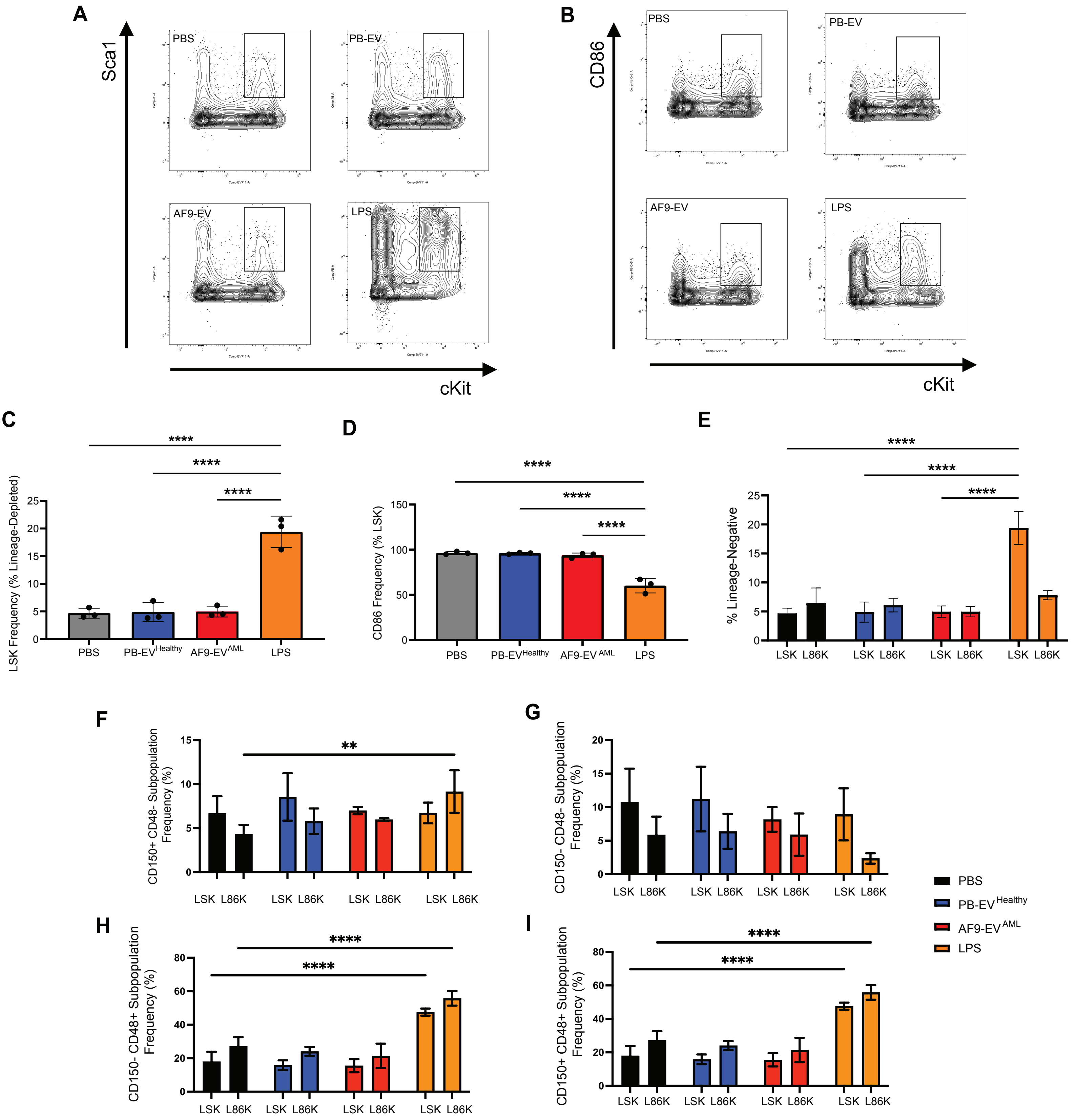


Supplementary Fig 6 – Analysis of LSK and L86K frequency following PBS, PB-EV$^{Healthy}$, AF9-EV$^{AML}$, and LPS challenge. Representative LSK (A) and L86K (B) flow cytometery gating. LSK Frequency (C), CD86 Frequency in LSK (D), comparative LSK and L86K frequency (E). HSPC subpopulation analysis in LSK and L86K fractions for: CD150+ CD48- population (F), CD150-CD48- population (G), CD150-CD48+ population (H), and CD150+CD48+ population (I).

#### Supplementary Table 1

List of real-time PCR primers sequences

| Gene | Species | Sequence Direction | Sequence |
|---|---|---|---|
| *Isg15* | Mus musculus | Forward | GGAACGAAAGGGGCCACAGCA |
| | Mus musculus | Reverse | CCTCCATGGGCCTTCCCTCGA |
| *Cxcl10* | Mus musculus | Forward | CCAAGTGCTGCCGTCATTTTC |
| | Mus musculus | Reverse | GGCTCGCAGGGATGATTTCAA |
| *Gapdh* | Mus musculus | Forward | AGGTCGGTGTGAACGGATTTG |
| | Mus musculus | Reverse | TGTAGACCATGTAGTTGAGGTCA |
| *IFN-alpha* | Mus musculus | Forward | CCTGAGAGAGAAGAAACACAGCC |
| | Mus musculus | Reverse | TCTGCTCTGACCACYTCCCAG |
| *Il-6* | Mus musculus | Forward | CCAAGAGGTGAGTGCTTCCC |
| | Mus musculus | Reverse | CTGTTGTTCAGACTCTCTCCCT |

**Supplementary Table 2**

List of anti-mouse antibodies used in immunofluorescence staining

| Marker | Clone | Manufacturer |
|---|---|---|
| CD117 (cKit) | 2B8 | BioLegend |
| Sca-1 | D7 | BioLegend |
| CD48 | HM48-1 | BioLegend |
| CD3a | 500A2 | BioLegend |
| CD11b | M1/70 | BioLegend |
| CD5 | 53-7.3 | BioLegend |
| CD4 | Gk1.5 | BioLegend |
| Gr-1 | R86-8C5 | BioLegend |
| B220 | RA3-6B2 | BioLegend |
| CD45.2 | 104 | BioLegend |
| CD45.1 | A20 | BioLegend |
| CD150 | TC15-12F12.2 | BioLegend |
| CD135 (FLK2) | A2F10 | BioLegend |
